# High-quality genome assembly of the endemic threatened White-bellied Sholakili *Sholicola albiventris* (Muscicapidae: Blanford, 1868) from the Shola Sky Islands, India

**DOI:** 10.1101/2024.08.20.608878

**Authors:** K L Vinay, Chiti Arvind, Naman Goyal, V.V. Robin

## Abstract

The White-bellied Sholakili (*Sholicola albiventris*) is an endemic, elevational restricted species occurring in the Shola Sky Islands of the Western Ghats of India. This unique understory bird, with a complex vocal repertoire, exhibits impacts of gene flow due to anthropogenic habitat fragmentation. Here, we present the first genome assembly for *Sholicola albiventris*, which was assembled using a combination of Nanopore and Illumina sequences. The final assembly is 1.083 Gbp, consisting of 975 scaffolds with an N50 of 68.64Mbp and L50 of 6. Our genome assembly’s completeness is supported by a high number of BUSCOs (99.9%) and a total of 4887 ultraconserved element (UCE) loci retrieved. We also report the complete mitochondrial genome comprising 13 protein-coding genes, 22 tRNAs, and 2 rRNAs. We identified 11.82% of the nuclear genome as repetitive and 36,000 putative genes, with 12017 genes functionally annotated. Our assembly showed a great synteny between *Taeniopygia guttata* and *Gallus gallus* chromosome level assemblies. This reference will be pivotal for investigating landscape connectivity, sub-population genetics, local adaptation, and conservation genetics of this high-elevation, range-restricted endemic bird species.

## Introduction

The White-bellied Sholakili (*Sholicola albiventris*) is a range-restricted, vulnerable passerine endemic to the high-elevation montane forests of the Shola Sky Islands of the Western Ghats of India. (Robin et al., 2017; *BirdLife International., 2024*) A phylogenetic study by Robin et al., 2017 revealed that the White-bellied Sholakili and one of its two congeneric sister species, the Nilgiri Sholakili (*Sholicola major*), are separated by an ancient biogeographic barrier within the Western Ghats. Formerly classified under the genera *Myiomela, Brachypteryx*, and *Callene* (Rasmussen & Anderton 2012; Dickinson & Christidis 2014; Rasmussen 2005), these species were determined to have a closer phylogenetic relationship within the taxa from the Himalayas and Southeast Asia, necessitating the establishment of a new genus. This slaty blue, monomorphic bird is often difficult to spot in the dense shola understory that it inhabits but is frequently heard due to its characteristic loud song. Notably, the White-bellied Sholakili possesses a highly complex birdsong, adding to its unique behavioral and ecological traits (Sawant et al., 2022).

The habitat of the White-bellied Sholakili originally comprised a bi-phasic mosaic of shola grasslands and shola forests. These shola ecosystems, characterized by patches of stunted tropical montane forests interspersed with open grasslands, create a unique landscape that supports a variety of endemic flora and fauna (Robin & Nandini, 2012). However, anthropogenic activities have significantly altered this landscape with increased invasive timber and agricultural lands over the past few decades (Arasumani et al., 2018). A study using microsatellite markers on the White-bellied Sholakili indicates a recent genetic differentiation in terms of shared alleles (DPS), which may have resulted from anthropogenic fragmentation (Robin et al., 2015). This genetic differentiation suggests that the population of the White-bellied Sholakili is isolated, resulting in reduced gene flow and loss of genetic diversity. Such genetic consequences can have long-term effects on the viability of the species, making conservation efforts even more difficult (Pavlova et al., 2017).

A high-quality reference genome for the White-bellied Sholakili will facilitate the conservation efforts and provide researchers with valuable resources for assessing population structure at a finer scale. This will enhance our understanding of how landscape modification affects species distribution and help uncover the regions of the genome involved in the complex song production. Here, we describe bShoAlb1.1, a de novo assembly constructed from a wild-caught White-bellied Sholakili. Using a hybrid assembly strategy with Nanopore Long Read technology and Illumina Short Read sequences, we have assembled the first published reference genome for the *Sholicola* genus.

## Methods

### Sample collection and DNA extraction

A female *S. albiventris* was captured using a mist-netting protocol (Robin et al., 2010). Blood was drawn from the ulnar vein and stored in the Queen’s lysis buffer. DNA extraction was performed using the Qiagen Blood and Tissue kit (Qiagen, Hilden, Germany) with slight modifications according to the manufacturer’s protocol. The concentration of the extracted DNA was measured using a Qubit 4 (Thermo Fisher Scientific Inc., USA) fluorometer, and its integrity was assessed on a 1% agarose gel. DNA was then sequenced on an R10 flowcell on the PromethION, targeting approximately 80x coverage for Oxford Nanopore long reads (ONT) and approximately 30x for 150 bp paired-end short reads on Illumina NovaSeq 6000.

### Read pre-processing

Oxford Nanopore reads underwent quality assessment using NanoPlot (Supplementary Figure S1), followed by any reminiscent adapter removal using Porechop v0.2.4 (De Coster & Rademakers, 2023; Wick et al., 2017). Adapter removed reads were subjected to quality trimming with Chopper v0.7.0, employing a quality threshold of Q>7 (De Coster & Rademakers, 2023). K-mer counts were then estimated using Meryl v1.4.1 (Rhie et al., 2020) with k=21, and GenomeScope2 was utilized for visualizing the generated histogram to determine genome size and heterozygosity (Ranallo-Benavidez et al., 2020). Illumina short reads quality was checked using Fastp v0.20.1 (Chen et al., 2018). Adapters and low-quality bases (Q<20) were trimmed using Trimmomatic v0.39 (Bolger et al., 2014).

### Nuclear Genome Assembly

Trimmed and quality-filtered long reads were used to de-novo assemble the genome using Flye v2.9.3-b1797 (--nano-hq, --asm_coverage 40, --genome_size 1.16g) (Kolmogorov et al., 2019). The ‘draft’ assembly was then subjected to contamination screening using the Foreign Contamination Screen (FCS-adapter and FCS-gx) suite (Astashyn et al., 2024) and found to have none. The draft assembly was then polished using both short reads and long reads. A total of five rounds of polishing was carried out. First, we used Medaka v1.11.3 (https://github.com/nanoporetech/medaka) to correct the assembly from Flye, which was then polished with one round of Racon v1.5.0 (Vaser et al., 2017). Three rounds of polishing with POLCA v4.1.10 (Zimin & Salzberg, 2020) were carried out using short reads to obtain the ‘polished’ assembly. We removed redundant haplotypes from the assembly using purge_haplotigs v1.1.3 (Roach et al., 2018) with coverage cutoffs of -l 5, -m 50, and -h 160. (Supplementary Figure S2). We further improved the assembly using ntLink with ntlink_rounds (w=250) (Coombe et al., 2021, 2023). Using the de-novo assembled mitogenome (see below), we removed any contigs associated with the mitogenome from the nuclear assembly. Further, reference-based pseudochromosome scaffolding was performed using RagTag v2.0.1. (Alonge et al., 2022) RagTag clusters, order, and orient assembly contigs based on a Minimap2 alignment of those contigs to a reference genome. We used the *Taeniopygia guttata* reference genome (GCA_003957565.4) to obtain the pseudochromosomes with default settings under the “scaffold” module within RagTag. We used gfastats v1.3.6 (Formenti et al., 2022) and compleasm v0.2.2 (Huang & Li, 2023) with the aves_odb10 dataset to evaluate the assembly quality and completeness after each step. We also assessed genome completeness by estimating the number of UCEs that could be retrieved from the genome. According to the online tutorial, we extracted the UCEs with Phyluce v1.7.3 (Faircloth, 2016) with 1000 bp flanking regions on both sides. Final scaffolds were renamed before uploading to NCBI.

### Mitochondrial genome assembly

The findMitoReference.py script within the MitoHiFi suite (Uliano-Silva et al., 2023) was employed to identify the closest available mitogenome to our species. Utilizing the Snowy-browed Flycatcher (*Ficedula hyperythra)* mitogenome (NC_058320.1) identified by findMitoReference.py as a reference, we assembled the mitochondrial genome from trimmed ONT reads using MitoHiFi v3.2.1 with default settings and annotated the assembly using MitoAnnotator v3.98 (Iwasaki et al., 2013; Sato et al., 2018; Zhu et al., 2023) (Supplementary Figure S3).

### Genome Synteny analysis

To evaluate the validity of our reference-guided scaffolding approach, we assessed genome synteny by aligning the Ragtag-scaffolded assembly with both the *Taeniopygia guttata* (GCA_003957565.4) and *Gallus gallus* (GCA_016699485.1) reference genomes. *Gallus gallus* was chosen as a reference due to its frequent use as a model organism for comparative studies. Utilizing the ‘nucmer’ module within MUMMER v4.0.0rc1, alignments were conducted and subsequently filtered using MUMMER’s delta_filter module, permitting many-to-many alignments with a minimum identity threshold of 70% (Marçais et.,al 2018). The resultant tab-delimited file of alignment coordinates was generated using the show_coords module and subsequently employed in OmicCircos package v1.40.0 (Hu et.,al 2014) in R v4.3.1 (R Core Team 2021) for Circos plot visualization of synteny.

### Repeat masking and Genome annotation

RepeatModeler v2.0.2 (Flynn et al., 2020), within Dfam Transposable Element Tools (TETools) container v1.88 (*Lerat et*.,*al 2017*) was employed (with - LTRStruct) to create a species-specific library of transposable elements and repetitive sequences of *Sholicola albiventris*. This species-specific library was merged with existing repeat libraries sourced from Dfam 0th and 3rd partitions (Storer et al., 2021) and RepBase Repeat Masker libraries (v20181026) (Bao et al., 2015; Jurka et al., 2005). The resulting combined library was then used in identifying and soft masking (-xsmall) the repeat elements using the RepeatMasker v4.1.2-p1 (Smit, AFA, Hubley, R & Green, P. *RepeatMasker Open-4*.*0*. 2013-2015 http://www.repeatmasker.org). We utilized BRAKER v3.03 to predict the gene models without the RNAseq data. BRAKER 3 predictions are based on successive training using GeneMark-EP+ and AUGUSTUS with extrinsic information of homologous protein sequences. To predict the locations of the genes, we employed the ProtHint pipeline within the BRAKER and trained with AUGUSTUS using vertebrate amino acid sequences from Vertebrata_OrthoDB_10 (Brůna et al., 2020, 2021; Buchfink et al., 2015; Gotoh, 2008; Hoff et al., 2019; Iwata & Gotoh, 2012; Lomsadze et al., 2005; Stanke et al., 2006, 2008). We ran InterProScan-5.66-98.0 (with AntiFam-7.0, CDD-3.20, Coils-2.2.1, FunFam-4.3.0, Gene3D-4.3.0, Hamap-2023_05, MobiDBLite-2.0, NCBIfam-13.0, PANTHER-18.0, Pfam-36.0, PIRSF-3.10, PIRSR-2023_05, PRINTS-42.0, ProSitePatterns-2023_05, ProSiteProfiles-2023_05, SFLD-4, SMART-9.0, SUPERFAMILY-1.75) (Jones et al., 2014) and eggNOG-mapper v2 (with eggNOG DB v5.0.2) on the protein domains identified by BRAKER (Cantalapiedra et al., 2021; Huerta-Cepas et al., 2019). Before the functional annotation, we sanitized the gff3 files using gfftk (https://github.com/nextgenusfs/gfftk). We then used funannotate v1.8.15 with the outputs from BRAKER, InterProScan 5, and eggNOG-mapper to annotate the genomes functionally. Statistics on the produced annotation have been generated using AGAT v1.0.0 (Dainat et., al 2020).

## Results and Discussion

### Genome assembly

Long read sequencing yielded 11.095 million reads (89GB) with a read length N50 of 10,371bp, longest read of 428.013Kbp (Supplementary Figure S1), and mean read quality of 15.4, leading to an estimated long-read depth of ∼77x. Additionally, short read sequencing yielded 221.97 million reads, and post quality and adapter trimming 221.95 million reads were retained, totaling 28.45GB with an estimated read depth of ∼25x for Illumina reads. GenomeScope2 estimated the genome size to be 1.16GB with a heterozygosity of 0.55% (Figure 1b).

**Figure 1.**
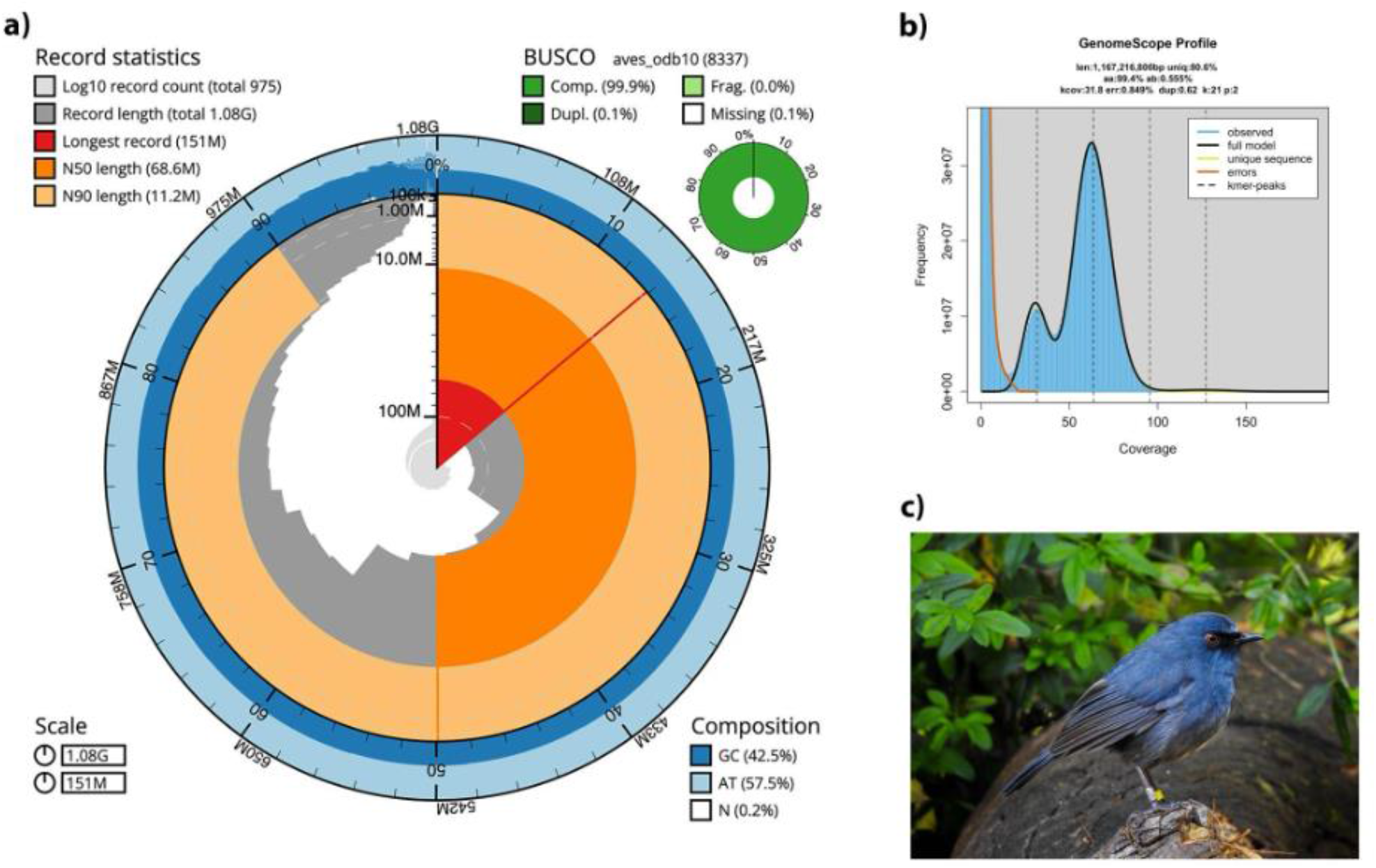
Genome characteristics of assembly bShoAlb1.1. a) BlobToolKit (Challis et., al 2020) snail plot showing a graphical representation of the quality metrics for the *S. albiventris* assembly (bShoAlb1.1). The circle plot represents the total size of the assembly. From the inside out, the central plot covers length-related metrics. The red line represents the size of the longest scaffold; all other scaffolds are arranged in size order, moving clockwise around the plot. Dark and light orange arcs show the scaffold N50 and scaffold N90 values. The dark versus light blue area around it shows mean, maximum, and minimum GC versus AT content. The BUSCO matrix is obtained from the compleasm. b) GenomeScope 2.0 profile of the k-mer spectra at k = 21 obtained using Meryl. The bimodal pattern observed corresponds to a diploid genome, and the k-mer profile matches that of low (<1%) heterozygosity. c) Photograph of a ringed *Sholicola albiventris* – Yellow - from Kodaikanal, India, picture credits: Vinay K L.

Our initial ‘draft’ assembly from Flye resulted in 2337 contigs, totaling 1.088 Gbp. After six rounds of long-read and short-read-based polishing, the number of contigs was reduced to 2036. Further, refinement through haplotig purging based on read coverage resulted in an improvement of contiguity and the number of contigs reduced to 1580, accompanied by a reduction in genome size to 1.081 Gbp and the contig N50 of 22.51Mbp. Minimizer graph-based scaffolding and orientation of contigs produced a total of 1254 scaffolds and 1293 contigs with a scaffold N50 of 33.76Mbp. Notably, fifty percent of the assembly was covered within the ten largest scaffolds (L50). The final assembly, integrated into pseudochromosomes, comprises 975 scaffolds with scaffold N50 of 68.64 Mbp, with the largest scaffold measuring 150.74 Mbp and an L50 of 6 scaffolds. See supplementary table S1 for a comparison of genome contiguity statistics of the recently published bird genomes.

Compleasm results showed that 99.81% (n = 8321) of the avian Benchmarking Universal Single-Copy Orthologs (BUSCO) were present in the final assembly. Detailed genome metrics and BUSCO scores are reported in Table 1. We recovered 4887 UCEs (96.9%, n = 5040), indicating the assembly’s high overall recovery and completeness, comparable to loci recovered from other Muscicapidae family genomes (Baudrin et al., 2023).

**Table 1.** Quality metrics for the assembly of *Sholicola albiventris* at various stages of the assembly pipeline with BUSCO scores

| Features | Draft assembly | Polished | Haplotigs purged | Scaffolded | Final (bShoAlb1.1) |
| --- | --- | --- | --- | --- | --- |
| # scaffolds | N/A | N/A | N/A | 1254 | 975 |
| Total scaffold length (Gbp) | N/A | N/A | N/A | 1.083 | 1.083 |
| Scaffold N50 (Mbp) | N/A | N/A | N/A | 33.76 | 68.64 |
| Scaffold L50 | N/A | N/A | N/A | 10 | 6 |
| Largest scaffold (Mbp) | N/A | N/A | N/A | 98.66 | 150.74 |
| # Gaps in scaffolds | N/A | N/A | N/A | 39 | 318 |
| # contigs | 2337 | 2036 | 1580 | 1293 | 1293 |
| Total contig length (Gbp) | 1.088 | 1.087 | 1.081 | 1.080 | 1.080 |
| Contig N50 (Mbp) | 22.52 | 22.51 | 22.51 | 26.82 | 26.82 |
| Contig L50 | 13 | 13 | 13 | 11 | 11 |
| Largest contig (Mbp) | 8.27 | 8.27 | 8.27 | 9.75 | 9.75 |
| aves_odb10 BUSCOs |  |  |  |  |  |
| Single | 99.69%,<br>8311 | 99.70%,<br>8312 | 99.80%,<br>8320 | 99.81%,<br>8321 | 99.81%,<br>8321 |
| Duplicated | 0.22%, 18 | 0.20%, 17 | 0.11%, 9 | 0.11%, 9 | 0.11%,<br>9 |
| Fragmented Class 1 | 0.02%, 2 | 0.02%, 2 | 0.02%, 2 | 0.01%, 1 | 0.01%, 1 |
| Fragmented Class 2 | 0.00%, 0 | 0.00%, 0 | 0.00%, 0 | 0.00%, 0 | 0.00%, 0 |
| Missing | 0.07%, 6 | 0.07%, 6 | 0.07%, 6 | 0.07%, 6 | 0.07%, 6 |
| Total | 8337 |  |  |  |  |

### Mitochondrial assembly

Our final complete circularized mitochondrial assembly obtained from MitoHiFi has a length of 16771bp (Supplementary Figure S3). The mitochondrial genome is similar to many other reported avian genomes with 13 protein-coding genes, 22 tRNAs, and 2 rRNAs with a GC content of 46% (Baudrin et al., 2023; Benham et al., 2023; Lan et al., 2024; Lu et al., 2019). Among the total annotated genes within the mitochondrial genome, 28 were on the heavy chain, and nine were on the light chain with a single non-coding control region (D-loop) of 1199bp length.

### Genome synteny

Our synteny analysis generally shows a large synteny between the pseudochromosomes of *Sholicola albiventris* and chromosomes of two other species (Figure 2). We recovered a high degree of one-to-one synteny between *Sholicola albiventris* and *Taeniopygia guttata*; however, it is worth noting that there was an absence of one-to-one synteny with the *Gallus gallus* assembly for the larger chromosomes (Figure 2). This indicates a likelihood of chromosomal splitting, attributed to the distinct phylogenetic relationship between taxa (Stiller et al., 2024) and varying numbers of chromosomes.

**Figure 2.**
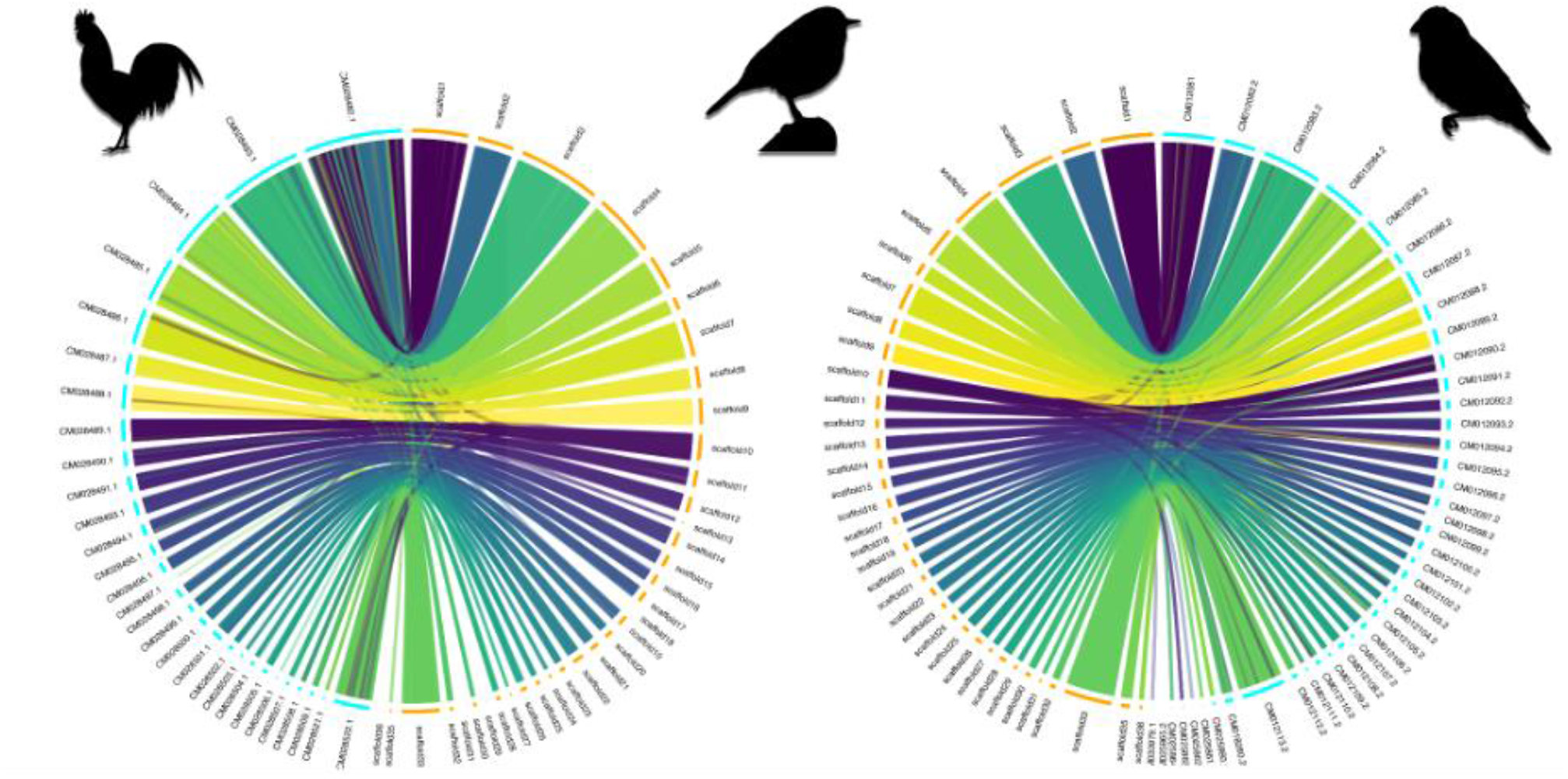
Circos synteny plots plotted using the OmicCircos package showing the comparison between the chromosomes of *Gallus gallus* (left) and *Taeniopygia guttata* (right) with pseudochromosomes of *Sholicola albiventris*. Chromosomes of the compared species are in the cyan-colored hemisphere, and S*holicola albiventris* is represented in the orange hemisphere. *Gallus gallus and Taeniopygia guttata* illustrations are reproduced from Phylopic (https://www.phylopic.org/).

Genome repeat content and annotation: RepeatMasker identified 11.82 % of the genome as repeat elements, of which 8.64% were interspersed repeats. Retroelements and DNA-transposons comprised 6.96% of repeats, and 1.69% of interspersed repeats were unclassified. Most of the remaining repeat elements were either simple repeats (1.76%) or satellites (0.93%). This is in line with the expected range of transposable elements for Aves (Sotero-Caio et al., 2017) and comparable to those reported in other Muscicapidae genomes (Baudrin et al., 2023; Peona et al., 2023). See Table 2 for a detailed classification of repeats. BRAKER initially found 36815 genes and 39052 mRNAs with a total gene length of 231827394bp. Genes compose 24.4% of the total genome, with a mean gene length of 6297bp. Functional annotation resulted in the names and/or descriptions assigned for 12017 genes and 13572 mRNAs. Additional annotation statistics and annotation files can be found in the Open Science Framework. (Vinay et al., 2024)

**Table 2:** Different classes of identified repeats within the bShoAlb1.1 genome.

|  | **number of**<br>**elements** | **length**<br>**occupied** | **Percentage of**<br>**sequences** |
| --- | --- | --- | --- |
| Retroelements | 207427 | 75147192 bp | 6.94% |
| SINEs: | 2173 | 271604 bp | 0.03% |
| Penelope | 0 | 0 bp | 0.00% |
| LINEs: | 132243 | 35251810 bp | 3.25% |
| CRE/SLACS | 0 | 0 bp | 0.00% |
| L2/CR1/Rex | 131762 | 35093953 bp | 3.24% |
| R1/LOA/Jockey | 0 | 0 bp | 0.00% |
| R2/R4/NeSL | 0 | 0 bp | 0.00% |
| RTE/Bov-B | 0 | 0 bp | 0.00% |
| L1/CIN4 | 481 | 157857 bp | 0.01% |
| LTR elements: | 73011 | 39623778 bp | 3.66% |
| BEL/Pao | 0 | 0 bp | 0.00% |
| Ty1/Copia | 0 | 0 bp | 0.00% |
| Gypsy/DIRS1 | 0 | 0 bp | 0.00% |
| Retroviral | 69498 | 37393552 bp | 3.45% |
| DNA transposons | 1267 | 166602 bp | 0.02% |
| hobo-Activator | 153 | 18244 bp | 0.00% |
| Tc1-IS630-Pogo | 100 | 15684 bp | 0.00% |
| En-Spm | 0 | 0 bp | 0.00% |
| MuDR-IS905 | 0 | 0 bp | 0.00% |
| PiggyBac | 0 | 0 bp | 0.00% |
| Tourist/Harbinger | 0 | 0 bp | 0.00% |
| Other (Mirage, P-element, Transib) | 0 | 0 bp | 0.00% |
| Rolling-circles | 2715 | 1547805 bp | 0.14% |
| Unclassified: | 50956 | 18305831 bp | 1.69% |
| Total interspersed repeats: | 93619625 bp | 8.64% |  |
| Small RNA: | 440 | 66018 bp | 0.01% |
| Satellites: | 4019 | 10046573 bp | 0.93% |
| Simple repeats: | 258494 | 19105522 bp | 1.76% |
| Low complexity: | 50504 | 3650726 bp | 0.34% |

Here, we present the first de novo assembled highly contiguous genome for the genus *Sholicola*, using a combination of Oxford Nanopore long reads and Illumina short read sequencing technologies. We believe this will serve as an essential resource for the investigations into landscape connectivity, sub-population genetics, local adaptation, and conservation genetics of this high-elevation, range-restricted endemic *Sholicola albiventris* and enhance our understanding of the genetic and evolutionary mechanisms underlying the unique characteristics and contribute towards the deeper understanding of the evolutionary trajectory of the avian genomes and ever-growing repository of avian reference genomes.

## Data Availability Statement

This Whole Genome Shotgun project has been deposited at GenBank under the accession JBDGPF000000000. The version described in this paper is version JBDGPF010000000. Raw reads with accession numbers SRR28564530 and SRR28558515 under the BioProject PRJNA1096119 are available from NCBI. Additional supporting data are available from the Open Science Framework. (Vinay et al., 2024) Associated scripts can be found in the GitHub repository (github.com/stachyris/ShoAlb_Ref_Genome).

## Acknowledgments

We thank the Bird Lab field team at IISER Tirupati for collecting the sample. The sample was collected with permits from the Tamil Nadu Forest Department (permit no WL5(A)/43781/2017). We are grateful to the Scientific Computing Facility at IISER Tirupati and members of the IT Department for HPC access. We thank Brant Faircloth for his valuable input during the project design and for providing computational resources. Portions of this research were conducted with high-performance computational resources provided by Louisiana State University (http://www.hpc.lsu.edu).

