## Supplementary for "High-quality genome assembly of the endemic threatened White-bellied Sholakili *Sholicola albiventris* (Muscicapidae: Blanford, 1868) from the Shola Sky Islands, India"

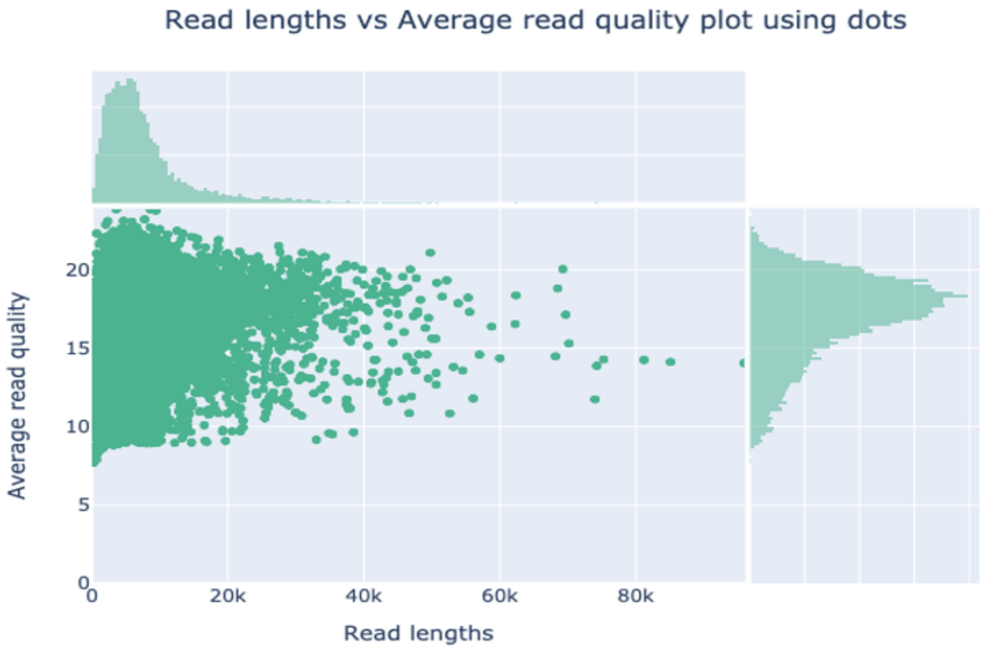


Figure S1: A dot plot generated by Nanoplot2 shows the distribution of read length and average read quality of raw reads.

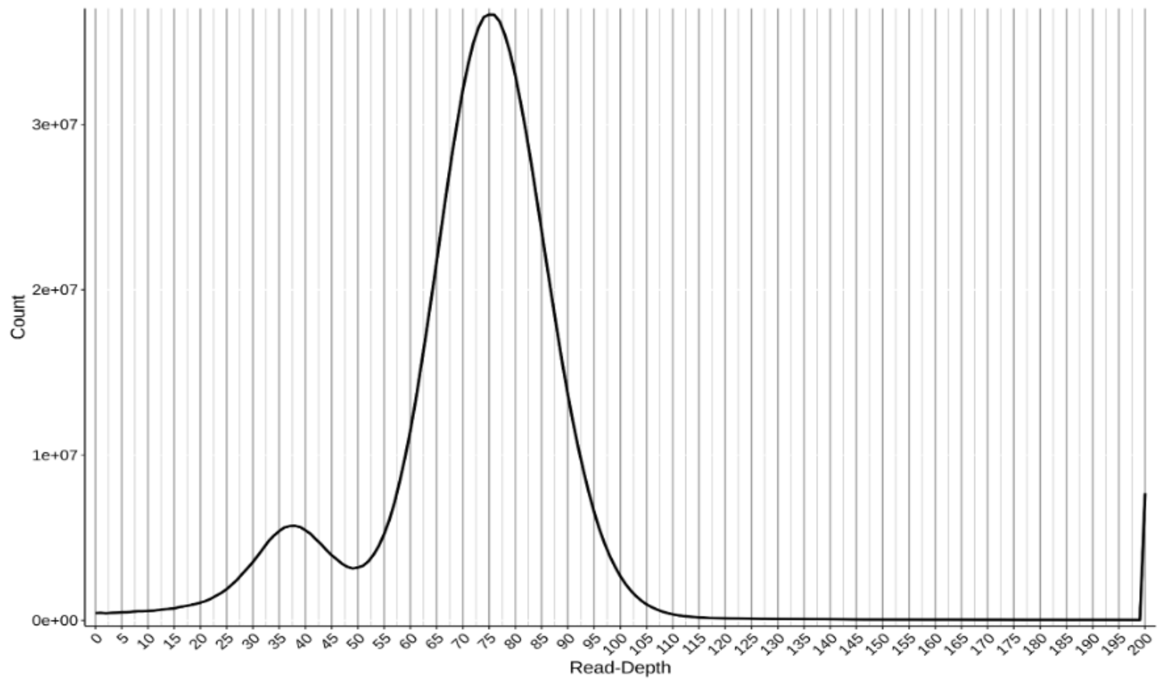


Figure S2: Coverage histogram generated by the 'hist' function of purge_haplotigs for the ONT reads showing the bimodal peak distribution. The peaks correspond to the coverage for the haploid and diploid genomes.

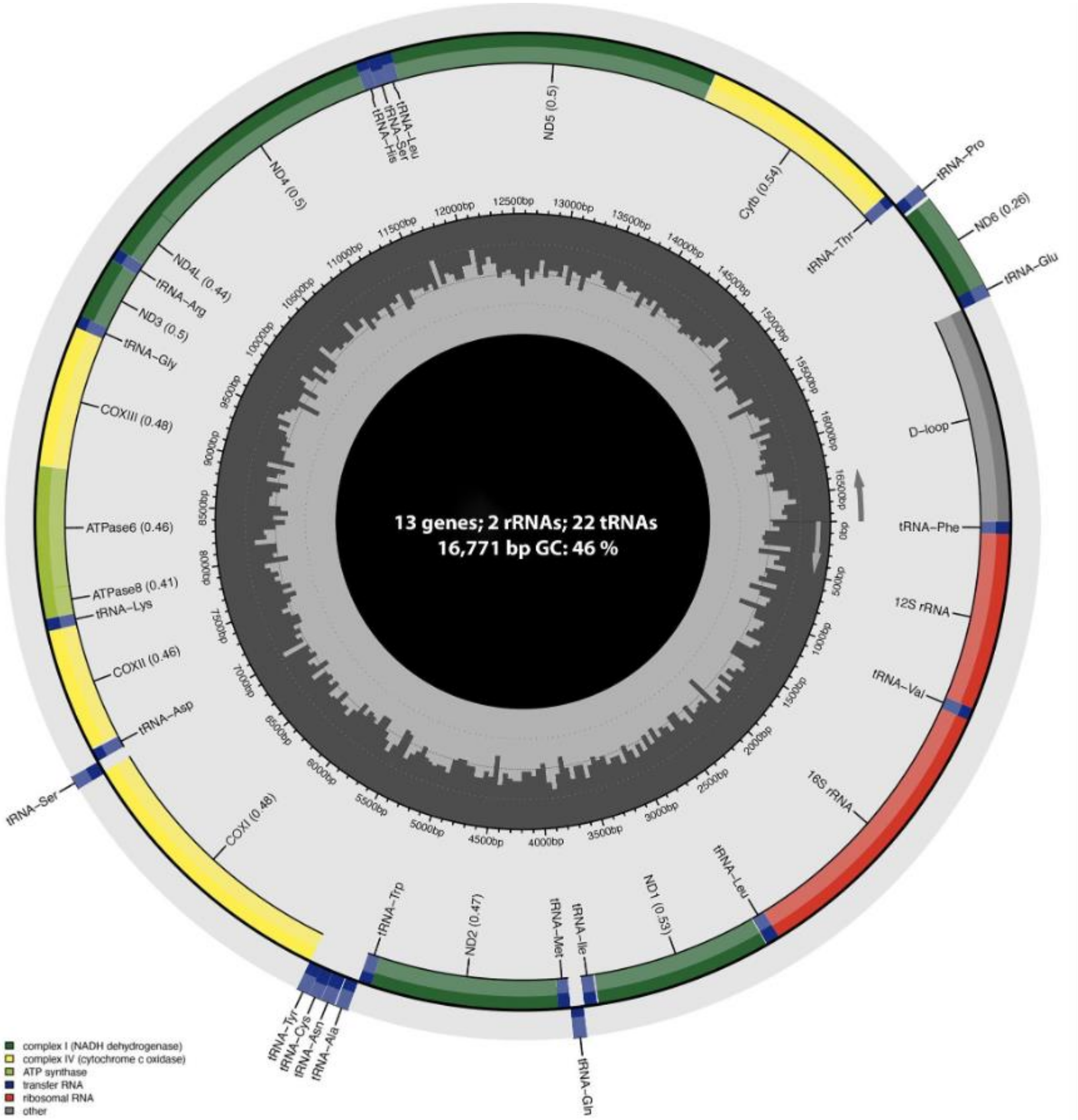


Figure S3: Circularized annotated mitochondrial genome of White-bellied Sholakili (*Sholicola albiventris)*. Figure generated by MitoAnnotator.

Table S1: Comparison of assembly statistics for the recently published bird genomes. Modified from Wiley et al. 2020

| Species | Reference | Method | Assembly size | # of contigs/ # of scaffolds | N50 contig | N50 scaffold | BUSCO complete % | % Repeats |
|---|---|---|---|---|---|---|---|---|
| *Sholicola albiventris* | This study | Nanopore (High Coverage)+Paired End | 1.08 Gb | 1293/975 | 26.82 Mb | 68.64 Mb | 99.81 | 11.87 |
| *Pavo cristatus* | Dhar et al., 2019 | Nanopore (low coverage) + short read | 932 Mb | 685,241/ 15,025 | 14.7 Kb | 0.23 Mb | Not reported | 7.3 |
| *Melospiza melodia* | Louha et al., 2019 | Paired end + HiC reads | 978 Mb | Not reported | 31.7 Kb | 5.6 Mb | 87.5 | 7.4 |
| *Rhegmatorhina melanosticta* | Coelho et al., 2019 | Paired end + 10× linked reads | 1.03 Gb | Not reported / 715 | 137 Kb | 3.3 Mb | 89.2 | Not reported |
| *Numida meleagris* | Vignal et al., 2019 | Paired end + Mate pair | 1.04 Gb | Not reported / 2,739 | 234 Kb | 7.8 Mb | 90.7 | 19.5 |
| *Colinus virginanus* | Salter et al., 2019 | Paired end + HiC reads | 866 Mb | Not reported /1,512 | Not reported | 66.8 Mb | 90.8 | Not reported |
| *Eremophila alpestris* | Mason et al., 2019 | Paired end + mate pair | 1.04 Gb | Not reported / 2,708 | Not reported | 10.6 Mb | 94.5 | Not reported |
| *Grus nigricollis* | Zhou et al., 2019 | Nanopore + paired end | 1.33 Gb | 1,837/ Not reported | 17.9 Mb | NA | 97.7 | 8.1 |
| *Melanerpes aurifrons* | Wiley et al., 2020 | Nanopore + paired end | 1.30 Gb | 441 / Not reported | 16.0 Mb | NA | 92.6 | 25.8 |
